# Exfoliated near infrared fluorescent CaCuSi_4_O_10_ nanosheets with ultra-high photostability and brightness for biological imaging

**DOI:** 10.1101/710384

**Authors:** Gabriele Selvaggio, Helen Preiß, Alexey Chizhik, Robert Nißler, Florian A. Mann, Zhiyi Lv, Tabea A. Oswald, Alexander Spreinat, Luise Erpenbeck, Jörg Großhans, Juan Pablo Giraldo, Sebastian Kruss

## Abstract

Imaging of complex (biological) samples in the near infrared (nIR) range of the spectrum is beneficial due to reduced light scattering, absorption, phototoxicity and autofluorescence. However, there are only few near infrared fluorescent materials known and suitable for biomedical applications. Here, we exfoliate the layered pigment CaCuSi_4_O_10_ (known as Egyptian Blue, EB) *via* facile tip sonication into nanosheets (EB-NS) with ultra-high nIR fluorescence stability and brightness. The size of EB-NS can be tailored by tip sonication to diameters < 20 nm and heights down to 1 nm. EB-NS fluoresce at 910 nm and the total fluorescence intensity scales with the number of Cu^2+^ ions that serve as luminescent centers. Furthermore, EB-NS display no bleaching and ultra-high brightness compared to other nIR fluorophores. The versatility of EB-NS is demonstrated by *in vivo* single-particle tracking and microrheology measurements in developing *Drosophila* embryos. Additionally, we show that EB-NS can be uptaken by plants and remotely detected in a low cost stand-off detection setup despite strong plant background fluorescence. In summary, EB-NS are a highly versatile, bright, photostable and biocompatible nIR fluorescent material that has the potential for a wide range of bioimaging applications both in animal and plant systems.

## Introduction

Fluorescence imaging provides important insights into the structure, function and dynamics of biological samples^1,2^. Imaging in the near infrared (nIR) spectral range (800-1700 nm) promises higher tissue penetration, higher contrast and lower phototoxicity due to reduced nIR light scattering and absorption^3–5^. However, these approaches are limited by the scarcity of nIR fluorescent materials. nIR fluorescent organic dyes such as indocyanine green bleach and are therefore not suitable for long term imaging^6,7^. In contrast, nanomaterials provide beneficial photophysical properties such as ultra-high photostability that would enable tracking in living systems without time constrains. nIR fluorescent nanomaterials include InAs quantum dots, lanthanide doped nanoparticles or semiconducting single-walled carbon nanotubes (SWCNTs)^8–12^. For example, SWCNTs have been used as building blocks for nIR imaging and as fluorescent sensors that detect small signaling molecules, proteins or lipids^13–19^. They can be chemically tailored and have been used to reveal spatiotemporal release patterns of neurotransmitters from single cells^2,15,20–22^. However, most nIR fluorescent nanomaterials often have low quantum yields, lack biocompatibility or are restricted to certain emission/excitation wavelengths. Therefore, there is a major need for novel nIR fluorescent and biocompatible nanomaterials for sophisticated applications such as long time single particle tracking in organisms or multiscale bioimaging such as stand-off detection in plants^23,24^.

One of the first colored pigments created by mankind is the calcium copper silicate called Egyptian Blue (CaCuSi_4_O_10_, EB), which has been synthesized and used as early as 2500 BC in Ancient Egypt^25^. Current ancient works of art decorated with EB have lost none of their vibrant color, a testimony to the remarkable chemical stability of this compound. Interestingly, bulk EB displays nIR fluorescence, which was only recently identified^26,27^ and attributed to a ^2^B_2g_-^2^B_1g_ electronic transition of the copper ion that ranges from 910 to 930 nm^26,28^. Bulk EB shows a remarkable high quantum yield of 10.5% for a nIR emitter compared to SWCNTs, quantum dots, metal nanoclusters and FDA-approved fluorophores like indocyanine green^5,28^. Recently, μm-sized monolayer sheets of EB were isolated by stirring in hot water for several days^29,30^. However, the remarkable properties of EB have not been explored for developing nIR luminescent nanomaterials for bioimaging applications. The layered structure of EB suggests that exfoliation procedures known from other 2D materials including graphene and transition metal chalcogenides could efficiently exfoliate it^31–34^. Such 2D materials have been shown to possess novel optoelectronic properties and are a rich playground for physics and chemistry^35^.

Herein, we use a facile tip sonication technique to exfoliate CaCuSi_4_O_10_ (EB) nanosheets (EB-NS) that allows one to control the nanomaterial size/thickness and retain the unique nIR fluorescent properties of macroscopic CaCuSi_4_O_10_. We report the photophysical properties of EB-NS and how nIR fluorescence scales with nanosheet size. Furthermore, we demonstrate the first use of this material for *in vivo* nIR microscopy and stand-off detection.

## Results and Discussion

The size of a nanomaterial determines its properties and interactions with the environment. For fluorescence imaging in cells or whole organisms fluorophores should be as small as possible to not perturb the system. Exfoliation into 2D sheets is one step but it is also important to decrease the lateral size into the nanoscale. Therefore, we exfoliated EB-NS *via* tip sonication, which allowed for the controlled decrease in height and diameter with sonication time (fig. 1a, fig. S1).

**Fig. 1:**
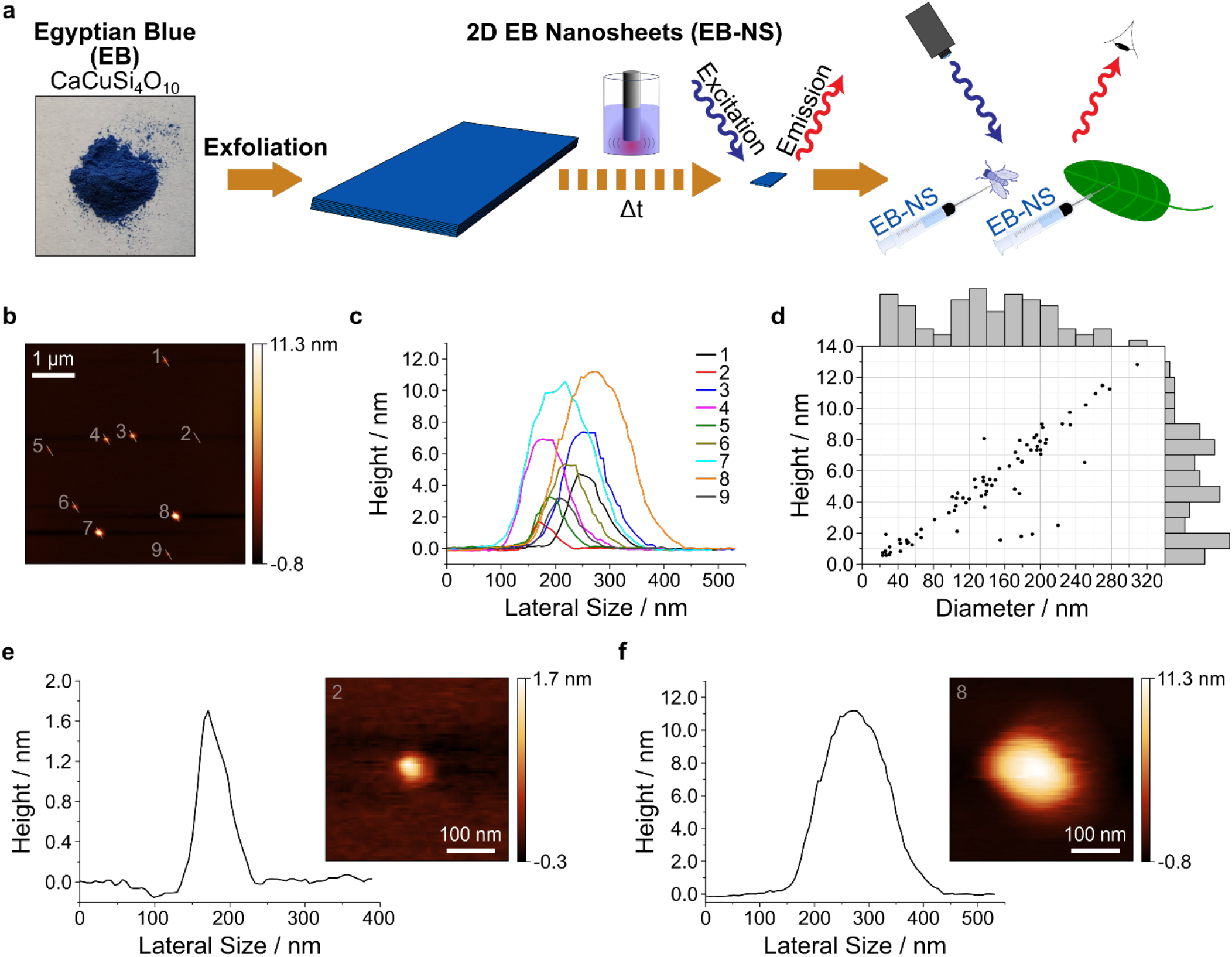
Exfoliation of Egyptian Blue (EB) into Egyptian Blue nanosheets (EB-NS). ***a** EB bulk powder (*CaCuSi_4_O_10_) *is exfoliated into EB-NS via tip sonication to study their near infrared (nIR) photophysical properties, and they are further used for in vivo nIR imaging in plants and Drosophila embryos. **b** Representative AFM image of EB-NS. **c** Corresponding height profiles (highlighted by a white line in b). **d** Height and diameter distribution after 6 h of tip sonication. The bars at the axis are histograms. n* = *78 EB-NS. **e,f** Height profiles and magnified images of the smallest and largest nanosheets shown in b.*

Tip sonication is a standard and widely used method to disperse nanomaterials such as carbon nanotubes^36^. 6 h tip sonicating in isopropanol yielded nanosheets of lateral sizes between 20-300 nm and heights of around 1-13 nm (fig. 1b-d). Atomic force microscopy (AFM) images (fig. 1b,c) indicate that EB height correlated linearly with diameter (fig. 1d). The height profiles of EB-NS (fig. 1e,f) typically contain a peak, which can be explained by the convolution through the AFM tip. The smallest observed height (≈ 1 nm) corresponds to the monolayer of EB bulk material reported in literature^29,37,38^ (fig. S2), but the exact height measured with AFM might vary depending on hydration and underlying substrate morphology^29^. To further purify EB-NS samples we typically used syringe cut-off filters (0.20 μm / 0.45 μm) to remove remaining larger particles from the samples.

In a next step, we explored how reducing the size and dimensionality of EB into the nanoscale regime affects its luminescence properties relative to macroscopic EB powder^27^. Fluorescence quantum yields of 1D materials such as SWCNTs have been shown to decrease with size, probably due to exciton diffusion and their collision with SWCNT ends^39^. Interestingly, EB fluorescence spectra did not change/shift with longer sonication times (fig. 2a) corresponding to smaller dimensions (fig. 1). EB-NS are extremely photostable as evidenced by the constant fluorescence emission intensity over several hours compared to the rapid bleaching of a typical organic dye (fig. 2b). These measurements were performed on an organic dye (Rhodamin B) and on EB-NSs, which were adsorbed on a glass surface under continuous excitation with a 561 nm laser (100 mW total power output) for > 2 h. Additionally, EB-NS show no change or shift in fluorescence emission in the presence of small redox active molecules that are known to affect the fluorescence of many dyes and fluorescent nanomaterials (fig. S3). EB-NS fluorescence is characterized by a large Stokes shift with a single emission maximum at ≈ 910 nm and an absorption maximum at ≈ 630 nm, as shown in the 2D excitation emission spectrum and in the reflectance spectrum (fig. 2c, fig. S4). EB-NS have a similar zeta potential (−22 mV) as spherical silica nanoparticles, which highlights that they can be dispersed and applied in aqueous solutions (fig. S5)^40^.

**Fig. 2:**
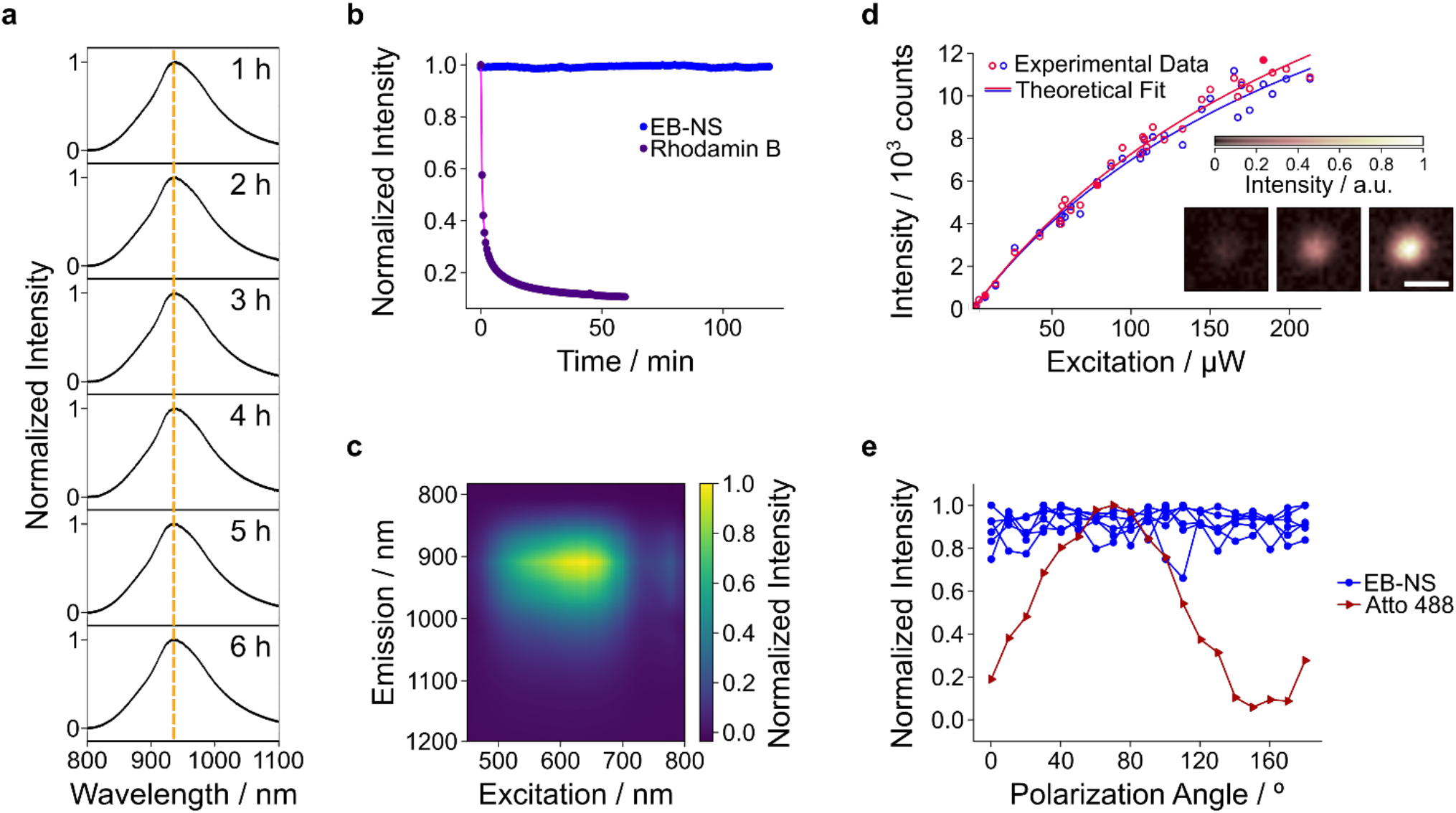
nIR fluorescence properties of EB-NS. **a** Normalized nIR fluorescence spectra of EB-NS after different sonication times show no wavelength shifts. **b** EB-NS fluorescence intensity (nIR) under continuous laser excitation (561 nm, 100 mW, corresponding to about 1 W/cm^2^) do not bleach. In contrast, the fluorescent dye Rhodamin B (in the visible) quickly bleaches. **c** 2D excitation-emission spectrum of EB-NS. **d** Fluorescence saturation measurements of a single EB-NS. Red circles show the data measured upon initial increase of excitation power, blue circles correspond to the subsequent decrease of the excitation power; red and blue lines represent the respective fits. The inset shows three images that were acquired at different excitation powers (corresponding data points are depicted with solid circles). Scale bar corresponds to 1 μm. **e** EB-NS display no fluorescence polarization. The dye Atto 488 is used as reference due to its distinct fluorescence polarization.

To estimate the number of luminescent centers in a single EB-NS, we performed single particle fluorescence saturation measurements of EB-NS using scanning confocal microscopy and pulsed laser excitation (20 MHz pulse rate). Only particles with sizes not exceeding 100 nm were studied (fig. 2d). Larger single sheets showed higher fluorescence intensities and could be distinguished from the smaller ones (see also fig. 4). This finding complies with the hypothesis that the number of luminescent centers is proportional to the size of EB-NS. To better understand how fluorescence scales with excitation intensity, the laser power was stepwise increased and decreased while fluorescence intensity was recorded. It was not possible to reach the emission saturation plateau with the maximum laser intensity. However, the slightly non-linear dependence between fluorescence intensity and laser power allowed us to fit the data with a fluorescence brightness saturation function *AI*_*exc*_ / (1+*I*_*exc*_/*I*_*sat*_), where *I*_*exc*_ is the excitation power, *I*_*sat*_ is the saturation power and *A* is a constant proportional to the absorption cross-section, quantum yield, microscope collection efficiency, etc^41^. The obtained saturation values (*I*_sat_) are 276 and 252 μW. The slightly lower saturation intensity value that was obtained for the process of diminishing excitation power could be caused by heating of the sample by the excitation light, which affects the fluorescence lifetime in bulk EB^28^.

The obtained saturation values correspond to a fluorescence emission of ≈ 13000 photons per laser pulse. With this number we estimated the number of luminescent centers in the order of 1300 by taking into account the average lifetime of the excited state (≈ 100 μs, fig. S6), the quantum yield (≈ 0.1^28^) of bulk EB, the quantum efficiency of the photodetector in this spectral region, the parameters of optical elements of the microscope and the acquisition time of the signal. From the crystal structure of EB it follows that there are 3.6 Cu^2+^ ions/nm^2^ of a single layer EB-NS^42^ (fig. S2). If every Cu^2+^ ion serves as luminescent center these numbers would be in agreement with a 36 nm large (squared) single layer EB-NS. From a resolution-limited EB-NS image we cannot derive the lateral size and height directly. Therefore, a 36 nm large monolayer EB-NS is the minimum size that is in agreement with the experiment. Nevertheless, these data suggest that a large proportion if not all Cu^2+^ ions act as luminescent centers. Furthermore, EB-NS showed no change of fluorescence intensity at different polarization directions (fig. 2e) compared to a typical organic dye (Atto 488). Similar to nIR emitting lanthanide complexes, bulk EB has a long fluorescence lifetime ranging from 107 μs to 142 μs at room temperature^28,43^ and this behavior is retained in EB-NS (fig. S6). The long fluorescence lifetime is most likely a consequence of the parity-forbidden nature of the transition, as the local D_4h_ symmetry of the Cu^2+^ ion in EB features an inversion centre^27^.

Larger EB-NS typically exhibit a layered structured as observed in scanning electron microscopy (SEM) images (fig. 3a,b, fig. S7). Exfoliation *via* tip sonication enabled us to continuously control the decrease in size of EB from large macroscopic particles to micrometer large structures down to EB-NS. These large and layered EB-NS can be imaged by a nIR fluorescence microscope (fig. 3c) but do not show a uniform fluorescence intensity profile, indicating that differences in thickness or geometry might affect fluorescence emission intensities. EB-NS smaller than the resolution limit of light microscopy can also be imaged in the nIR (fig. 3d,e), which proves that the nIR fluorescence properties of macroscopic EB particles are retained in nanosheets. These observations raise the question of how fluorescence intensity scales with size and if there is a lower limit, which is also relevant for potential bioimaging applications. Unfortunately, nIR microscopy is not able to directly assess the EB-NS size below the resolution limit. The resolution limit of light microscopy is described by the Abbe law (λ/2 ≈ 910 nm/2 ≈ 450 nm) and such resolution-limited single EB-NS are shown in fig. 3e. The photon counting experiments from fig. 2d show that there are many luminescent Cu^2+^ centers in one EB-NS, but because of the resolution limit it is not possible to assess the actual size. To provide an unambiguous answer, a correlative method that measures both size and fluorescence intensity of the same single nanosheet is required.

**Fig. 3:**
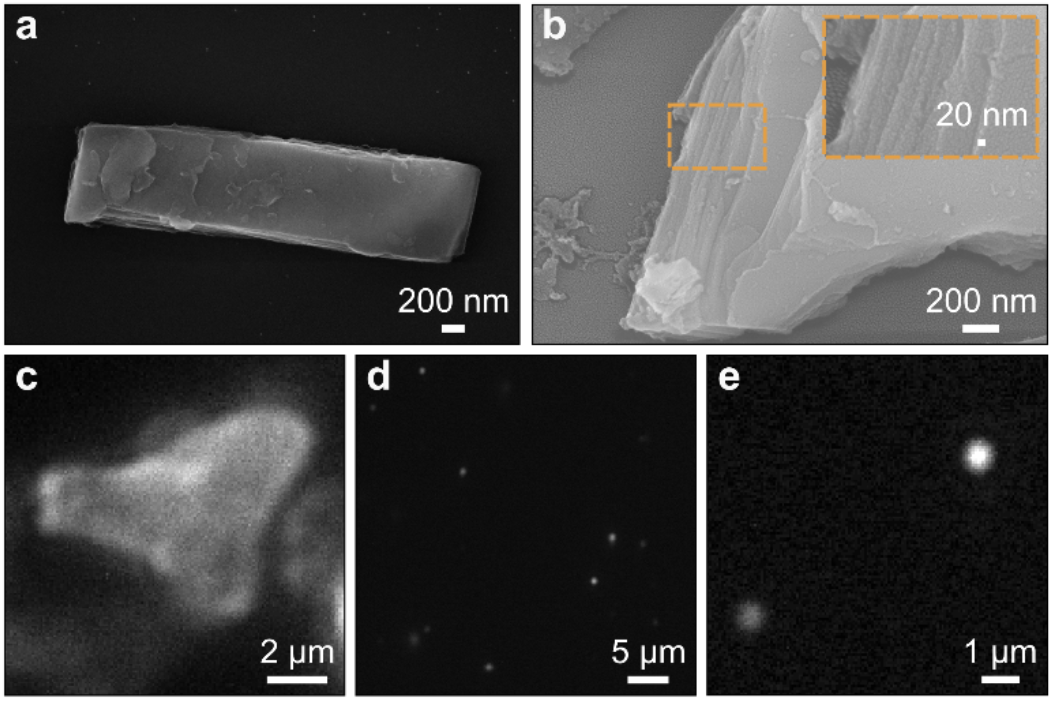
nIR imaging of EB-NS. **a,b** Scanning electron microscopy (SEM) images of larger EB particles. **c** nIR image of a larger EB-NS. **d,e** nIR images of EB-NS at increasing magnification show that single EB-NS can be resolved via nIR fluorescence microscopy down to the resolution limit of light microscopy. In e, EB-NS with a diameter below the resolution limit (≈ 500 nm) are shown. Excitation wavelength 561 nm.

To understand if and how nanomaterial fluorescence intensity is affected by size, we performed single particle tracking of EB-NS in a viscous glycerol solution. This approach allowed us to simultaneously quantify fluorescence intensity and Brownian motion as a measure of size of the same EB-NS. We used the maximum intensity during the whole trace as a measure of fluorescence intensity to account for out of focus movement or rotations. A size equivalent can be derived from the trace of the random walk (Brownian motion) by employing the Stokes-Einstein equation. In fig. 4a-c, nIR fluorescence images of EB-NS are shown (inverted color). The EB-NS in fig. 4a is larger than the resolution limit, while the EB-NS in fig. 4b,c are smaller. The EB-NS in fig. 4c can barely be seen with the same contrast settings as in fig. 4a,b. However, if the contrast is changed, the EB-NS becomes visible (fig. 4d). The trajectories (fig. 4e) show that the brighter particles move slower, indicating again that fluorescence intensity depends on size. To further investigate this finding, mean square displacement plots (MSD, 〈*r*^2^〉) were determined. The MSD plots in fig. 4f corresponding to the trajectories in fig. 4e show slightly superdiffusive behavior, possibly because the trajectories were not corrected for drift. To estimate the EB-NS size, we determined the MSD for a certain lag time *τ* and calculated the diffusion constant 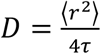. The Stokes radius *R* is related to the diffusion constant *via* the Stokes-Einstein equation, 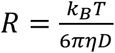 where *η* is the dynamic viscosity of the solvent, *T* the temperature and *k*_B_ the Boltzmann constant. As EB-NS are rather anisotropic with a large aspect ratio (fig. 1, fig. 3, fig. S7), we assumed that the diffusion is dominated by the diameter and not the height similar to a spherical particle. Approximations that correct the Stokes-Einstein equation for anisotropy have been used for example to analyze carbon nanotube diffusion and length, but there is no analytical solution available for 2D nanosheets^44^.

**Fig. 4.**
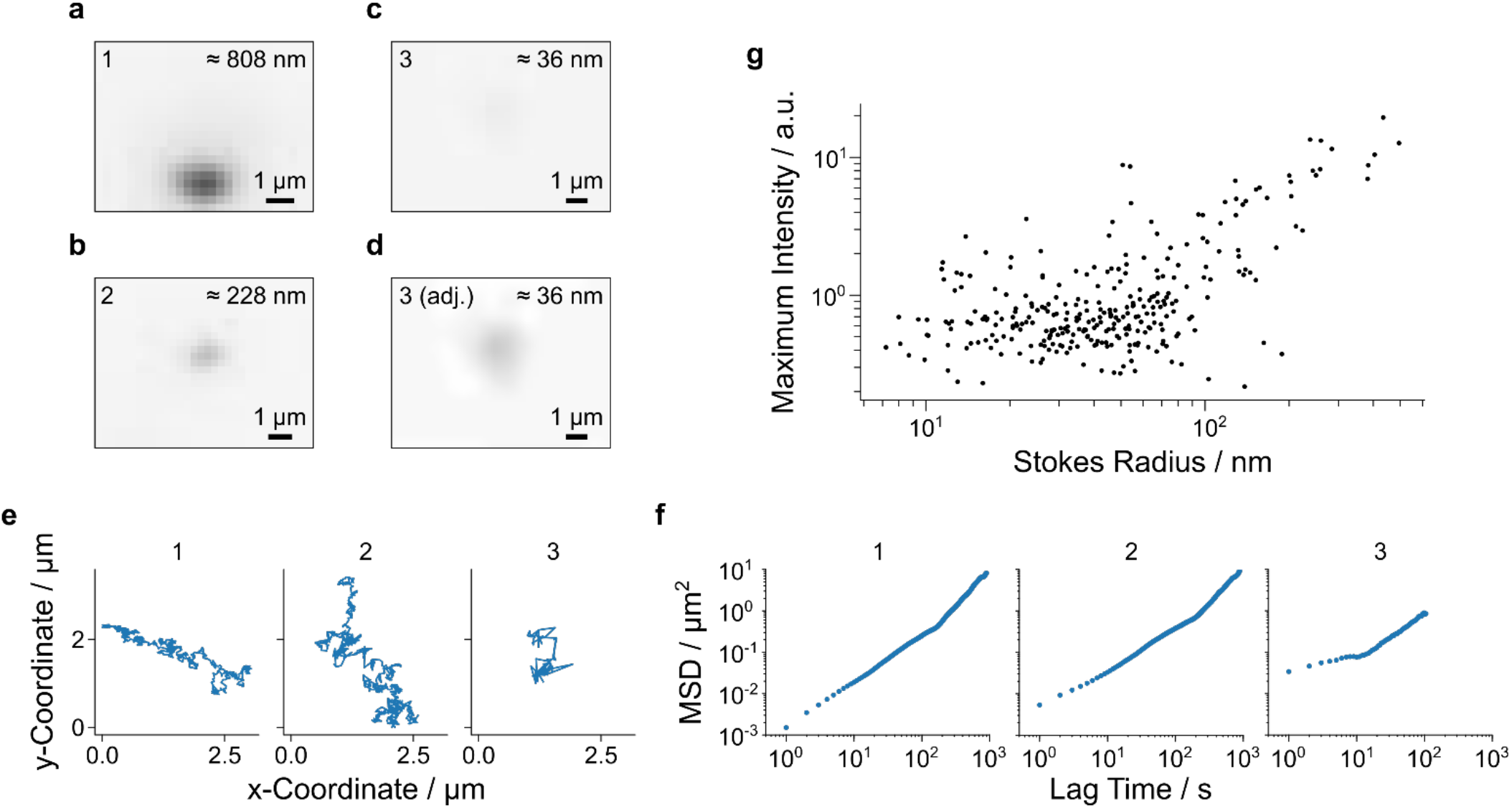
Correlative measurement of size and fluorescence intensity of EB-NS. Single EB-NS were imaged and their fluorescence intensity quantified in a viscous glycerol solution to slow down diffusion. Additionally, Brownian motion of the nanosheet was analyzed to derive a hydrodynamic radius via MSD analysis and the Stokes-Einstein equation. Therefore, both fluorescence intensity and hydrodynamic radius of single EB-NS are accessible. **a** EB-NS with a size above the resolution limit. **b,c** EB-NS with a size below the resolution limit. **d** EB-NS shown in c with adjusted contrast. The images of the three nanosheets 1-3 shown in a-d are presented in inverted colors and the estimated hydrodynamic diameters are displayed in each top right corner. **e** Trajectories corresponding to the EB-NS 1-3 from a-c. **f** MSD plots of EB-NS 1-3 corresponding to the trajectories in e. **g** Fluorescence intensity as a function of the Stokes radius of EB-NS of different sizes. Even the smallest EB-NS (< 10 nm) still fluoresce. The general trend indicates that EB-NS of larger size are brighter. N = 4 independent samples, n = 297 EB-NS.

The fluorescence intensity of EB-NS of different size as a function of Stokes radii is shown in fig. 4g. We calculated radii from 7 to 494 nm, with most sheets having a Stokes radius of 10 to 100 nm, similar to the AFM measurements shown in fig. 1b. The predicted Stokes diameters for the sheets in fig. 4a-c are 808 nm for “EB-NS 1”, 228 nm for “EB-NS 2” and 36 nm for “EB-NS 3”. These results show again that larger EB-NS are brighter than smaller ones, indicating that fluorescence intensity correlates with the number of luminescent Cu^2+^ centers and thus the volume. The spread of data points along the fluorescence intensity axis at a given Stokes radius is most likely due to different layer numbers for sheets of similar diameter. However, purification approaches that lead to samples of precise layer number and sheet diameter could further enhance our understanding of how EB-NS fluorescence scales with the dimensions. Together with the photon counting experiments in fig. 2, these results indicate that even the smallest EB-NS are fluorescent and fluorescence emission intensity scales with the number of Cu^2+^ ions. Therefore, dimensionality does not appear to affect fluorescence properties beyond the change in the number of luminescent Cu^2+^ centers.

To demonstrate the potential of EB-NS for bioimaging applications, we performed single-particle tracking and microrheology measurements of EB-NS in embryos of the fruit fly *Drosophila melanogaster*. This fly species is a widely employed model organism for studies ranging from fundamental genetics to developmental cell biology. During embryonal development the nuclei arrange in complex patterns mediated by microtubules and the actin cortex, but the underlying mechanisms and mechanics are poorly understood^45^. Following fertilization, the embryo develops as a syncytium^45^ in which the nuclei arrange in a regular two-dimensional array linked to the actin cortex of the plasma membrane (fig. 5a).

**Fig. 5:**
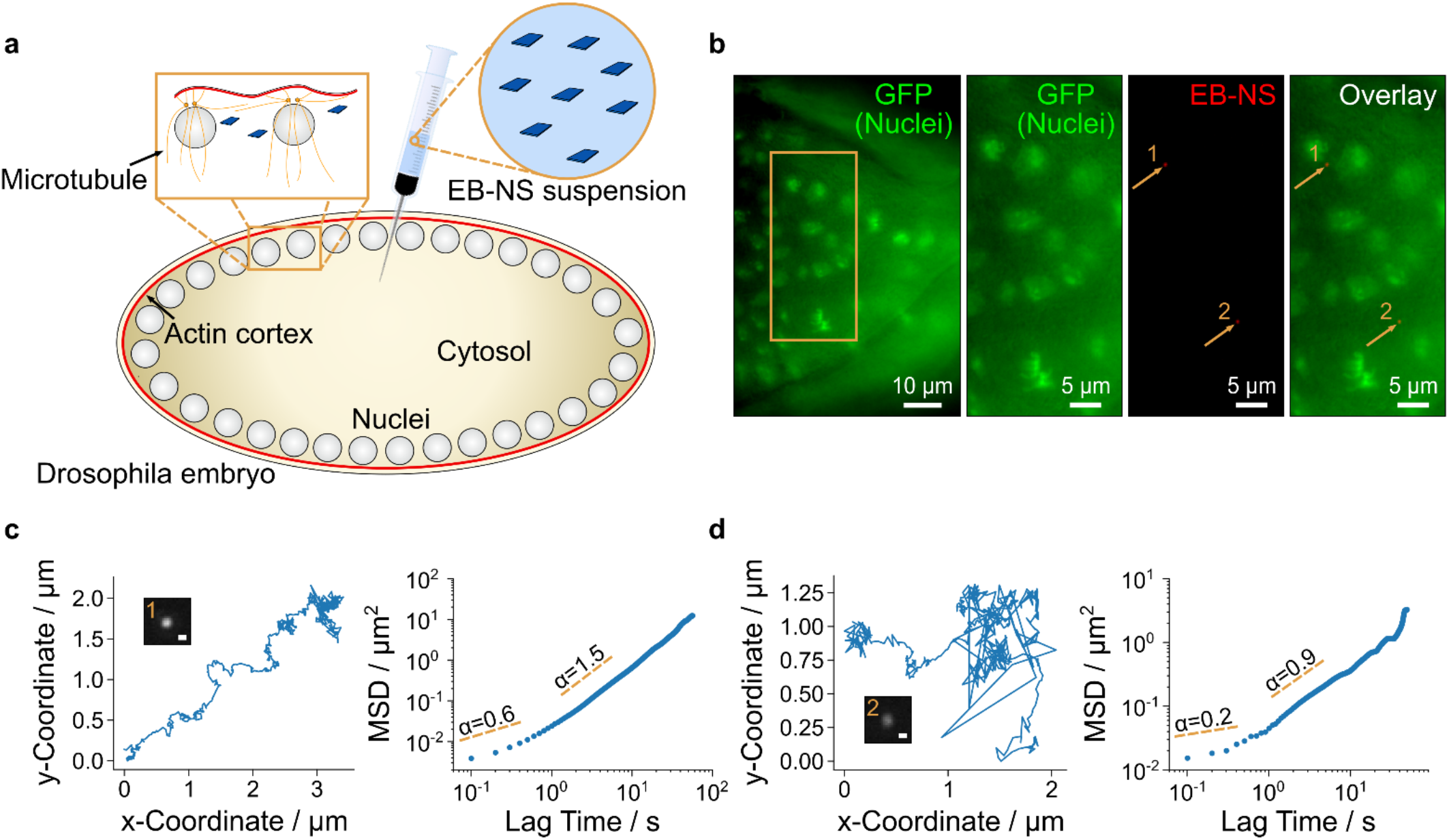
*In vivo* single-particle tracking and microrheology with EB-NS. **a** Experimental scheme for microinjection of EB-NS into syncytial Drosophila embryos. During the syncytial blastoderm stage, the peripheral nuclei are linked to the actin cortex of the plasma membrane (red). **b** EB-NS (red) inside a Drosophila embryo during gastrulation stage (stage 6) expressing Histone 2Av-GFP (green) that labels the nuclei. Overview (left image), magnified region (orange rectangle) with GFP (green), nIR EB-NS (red) and overlay channels are shown. Arrows pinpoint to the 2 EB-NS in this vield of view. **c,d** Traces (left plot) and MSD (right plot) of EB-NS 1 and 2 shown in b. The slopes (extracted from linear fits of two different lag time regions) indicate a subdiffusive behavior at short time scales and active motion (superdiffusion) at longer time scales. Scale bars in EB-NS images correspond to 0.5 μm.

The dynamics of the nuclei and their associated centrosomes is determined by the cortical link and internuclear interactions mediated by microtubules^46^. It has been hypothesized that the viscoelastic properties and the molecular motors play a key role in this process^45^. To elucidate the micromechanical properties of the cellular matrix, we introduced EB-NS into *Drosophila* embryos and tracked single EB-NS *via* nIR fluorescence microscopy. Imaging in *Drosophila* is challenging due to autofluorescence in the visible region and its high sensitivity to phototoxicity, therefore nIR imaging would be very beneficial.

For assessing the relative location of the EB-NS in *Drosophila*, mutants expressing GFP-labelled histones in their nuclei were used (fig. 5b). Using this approach, we followed the traces of single EB-NS *in vivo* (fig. 5b), allowing long-term imaging without bleaching or developmental abnormalities despite the high temporal resolution with frame rates > 10 Hz. In addition, the low autofluorescence in this spectral range enabled a higher contrast of the nIR EB-NS image compared to the visible GFP image (fig. 5b). These results highlight the great advantages of a non-bleaching nIR nanoscale fluorophore compared to typical organic dyes. The observed traces of EB-NS (fig. 5c,d) depend on the viscoelastic environment and active processes such as flow or contractions inside the embryo and thus provide important insights. The MSD (〈*r*^2^〉) is related to the diffusion constant by 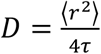. For the EB-NS tracks shown in fig. 5c,d the diffusion constant *D* is 3.1×10^−14^ m^2^ s^−1^ and 9.5×10^−15^ m^2^ s^−1^ (linear fit of the MSD for the linear regime < 30 s). These diffusion constants correspond to dynamic viscosities (*η*) of 14 mPa s and 49 mPa s, derived by using the Stokes-Einstein equation. These data show that there are different viscoelastic environments in the embryo that can be mapped with EB-NS. The measured values are higher than those of water (1 mPa s) but lower than reported values (up to 1 Pa s)^45^. These differences can be attributed to the size of EB-NS, which is much smaller than the typical micrometer-sized probes used for microrheology. EB-NS are likely to probe the mesh-size of the embryo’s dense cytosol on the nanoscale without being easily trapped in between cellular filaments and other subcellular barriers. Therefore, EB-NS probe mechanical properties on a size scale that is not accessible by typical micrometer-sized beads used for microrheology^45^. Active processes and interactions inside the embryo (including movement of the nuclei, contractions and even flow) increase the movement of the nanoprobes beyond diffusion, which can be seen from the increasing slope in the MSD plots (fig. 5c,d) ^47^. This behavior (superdiffusion) is indicative of active forces such as contractions inside the embryo on longer time scales. The MSD plots of the EB-NS closer to nuclei (EB-NS 1 in fig. 5) differ from the ones in the cytosol in between (EB-NS 2 in fig. 5.). The higher MSD slope of the EB-NS close to the nucleus indicates that there are more active forces beyond diffusion involved. A possible explanation are forces exerted by the microtubules (in the spindle close to the nuclei) that move the nuclei apart from each other. Our results show that EB-NS are powerful probes for such studies. In this context, nanomaterials in general can become important tools to study viscoelastic properties of (living) matter. For example nIR fluorescent carbon nanotubes have been recently used to explore the extracellular space between neurons^48^.

In the context of biological applications, toxicity is a major concern and for example a potential drawback of quantum dots that contain toxic elements and 2D materials in general^49,50^. To evaluate cytotoxicity, viability assays with EB-NS exposure to different cell lines (A549, 3T3 and MDCK-II) were performed. We observed no significant effects on the viability of these cell lines, highlighting the biocompatibility of EB-NS (fig. S8). In contrast, Cu^2+^ from soluble *CuSO_4_* decreased cell viability, which shows that EB-NS does not release relevant Cu^2+^ ion concentrations as expected from a stable silicate. Overall, this study demonstrates that EB-NS are powerful and biocompatible nanoprobes in living organisms for single-particle tracking and microrheology measurements in the transparent nIR window.

Another potential application of nIR fluorescent EB-NS is non-contact stand-off detection of nanoprobes in living organisms with minimal perturbation. The low autofluorescence in the nIR region provides a high contrast, but stand-off detection requires especially bright fluorophores because only a small portion of emitted light reaches the detector. Recently, nIR fluorescent SWCNT sensors enabled plants to report to electronic devices chemical signaling processes, pollutants in the environment^51^ and explosives^52^. Optical nanosensors are poised to allow the engineering of smart plants that communicate with and actuate agricultural and phenotyping devices for improving crop productivity and resource use^24^.

Herein, we developed a low cost optical setup to image the nIR emission of EB-NS from a distance > 10 cm with unprecedented brightness compared to several other nIR fluorophores. The setup consisted of a white light LED and a camera equipped with nIR (900-920 nm) longpass filters (fig. 6a). The high brightness of EB-NS allowed us to use a simple CMOS-camera (Si-based) instead of expensive electrically or nitrogen cooled InGaAs cameras (> 40k €), which would have a much higher quantum yield in this spectral range. In the visible image (fig. 6b, top) one can easily distinguish SWCNTs (black) from ICG (green), whereas the solution with EB-NS is transparent similar to the water control. In contrast, in the corresponding nIR fluorescence image (fig. 6b, bottom), EB-NS are significantly 2x brighter than ICG and 10x brighter than SWCNTs (fig. 6b, fig. S9). Interestingly, stand-off detection of the nIR emission of EB-NS is also possible without LED excitation: room or sun light alone is sufficient to generate detectable nIR fluorescence from EB-NS (fig. 6e).

**Fig. 6:**
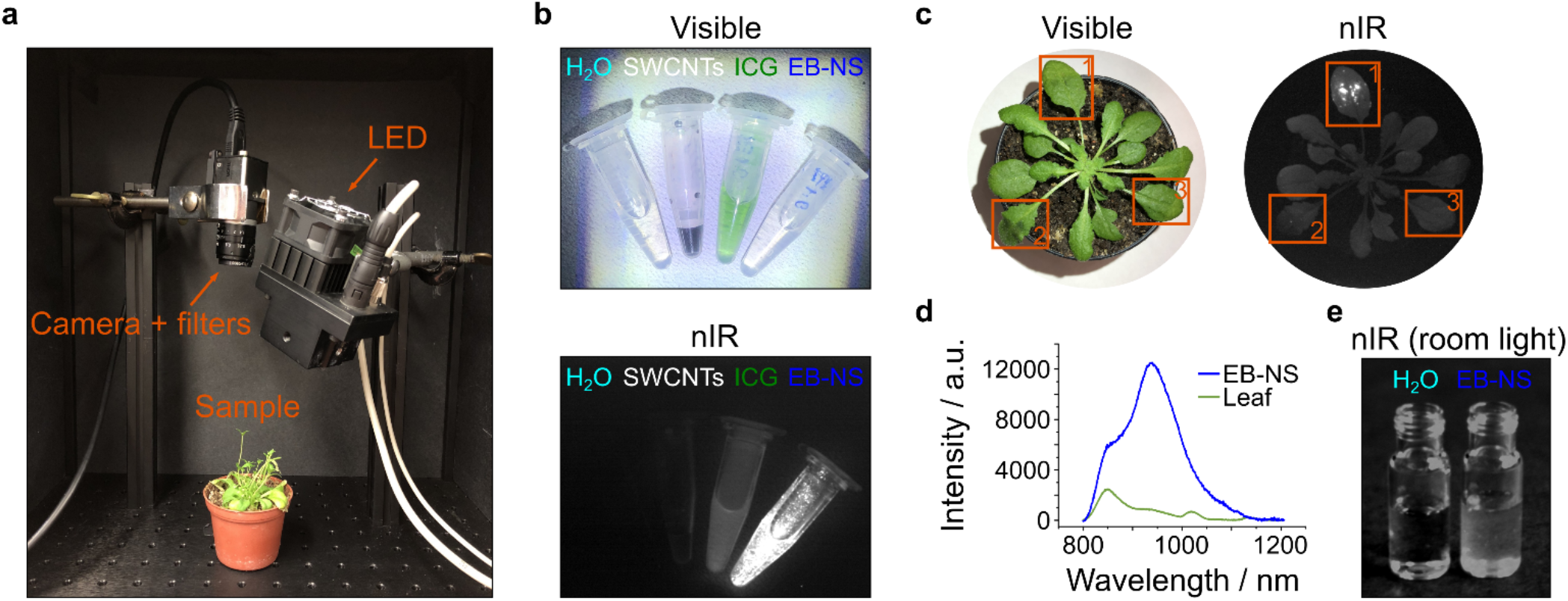
Stand-off detection of EB-NS fluorescence in plants through low cost and widely available imaging devices. **a** Picture of a plant (Arabidopsis thaliana) placed in a low cost stand-off imaging system, which consisted of a LED, nIR filters and a Si-CMOS camera. **b** Visible (top) and nIR fluorescence (bottom) images of EB-NS (≈ 0.1 mg/mL) compared to other nIR nanomaterials and fluorophores at similar concentration (single-walled carbon nanotubes (SWCNTs), indocyanine green (ICG)). Water is used as negative control. **c** Visible (left) and nIR (right) images of an Arabidopsis plant, which was infused with EB-NS (frame 1), SWCNTs (frame 2) and buffer only (frame 3). **d** The nIR fluorescence spectrum of the leaf confirms the presence of EB-NS and its strong emission compared to the leaf background. Both results demonstrate the unprecedented brightness of EB-NS compared to state-of-the-art nIR nanomaterials (SWCNT) and that this platform can be applied for stand-off detection using a low cost optical setup. **e** The EB-NS emission can even be detected without LED excitation under room light conditions.

Buffered EB-NS were infused into the leaves of *Arabidopsis thaliana*, a well-established plant model system (fig. 6c,d)^51^, using a standard method for nanomaterial delivery into plant leaves *in vivo*^51^. The same approach was used for the infiltration of SWCNTs in separate leaves in the same plant, which served as a visual comparison with EB-NS. As it can be observed from the nIR image in fig. 6c, plant leaves are characterized by a background autofluorescence in this wavelength region that hinders the visualization of SWCNTs fluorescence (frame 2) through a Si-based CMOS camera, whereas EB-NS can be detected with a strong fluorescence signal (frame 1). Similarly, fluorescence spectra of these leaves (fig. 6d) show the strong EB-NS emission compared to the (autofluorescence) background. These stand-off detection experiments highlight that EB-NS can be delivered into living organisms with high background fluorescence such as plants and can be detected using low cost and widely available stand-off imaging devices.

Our results show that EB-NS have extremely useful properties for nIR bioimaging applications including ultra-high photostability, extreme brightness for nIR fluorophores and biocompatibility. The remarkable properties of EB-NS are retained even when the dimensions of the nanosheets are reduced. Our data indicate that most if not all copper ions serve as luminescent centers that emit nIR light. Therefore, size reduction does not decrease the nIR fluorescence quantum yield. It only affects the absorption cross section by changing the number of light absorbing Cu^2+^ ions. In contrast, the quantum yield of nIR fluorescent SWCNTs increases with length. As a consequence, shorter SWCNTs (< 100 nm) are much less bright than longer ones^39^. Therefore, SWCNTs cannot be shortened without disadvantages. Although it has been reported that for bioimaging applications the nIR-II (1000-1700 nm) windows further improves tissue penetration^5^, a major advantage of the EB-NS fluorescence in the nIR-I region (800-1000 nm) is the wide availability of cameras^53,54^. Instead of expensive (>40k €), liquid nitrogen-cooled low resolution InGaAs-based cameras, low cost (1k €) high-resolution Si-based cameras can be used for detection. In the past, (6,4)-SWCNTs with smaller diameter that emit in the nIR-I region at 870 nm have been isolated exactly for this purpose^55^. Even though the quantum yields of Si-based cameras rapidly decrease in the nIR (around 5% > 900 nm), they are still able to image even single EB-NS using a normal microscope or a low cost stand-off detection device. Furthermore, there exist (bulk) pigment homologues (Ba, Sr), which have a red-shifted emission spectrum up to 1000 nm. Nanosheets of those materials could therefore extend the spectral range further into the nIR. The capability of imaging EB-NS even with only room light excitation using Si-based cameras is particularly relevant for commercial applications beyond the well-equipped research laboratories or biomedical research, *e.g.* the engineering of smart plants for high-throughput phenotyping and precision agriculture. Another major advantage of EB-NS is their low toxicity, proven by the performed viability assays on cells (fig. S8). With our study we add EB-NS to the library of 2D materials in general and especially exfoliated silicates and clays^56,57^. In the future, EB-NS and their physicochemical properties could be further investigated to find non-expected properties as in other 2D materials such as transition metal dichalcogenide nanosheets^58^. Additionally, the surface chemistry could be tailored with biomolecules similar to silica nanoparticles and silicate coated core-shell nanoparticles^59,60^.

## Conclusion

In summary, we developed a method to exfoliate the calcium copper silicate Egyptian Blue into nanosheets (EB-NS) and demonstrated that their nIR optical properties are retained at the nanoscale with copper ions as luminescent centers. We add a novel 2D material to the class of nIR fluorophores with ultra-high photostability, brightness and biocompatibility. EB-NS have a large potential for demanding biological applications such as *in vivo* imaging in complex biological matrices, including animal and plant model systems.

## MATERIALS AND METHODS

### Exfoliation of Egyptian Blue (EB) into Egyptian Blue Nanosheets (EB-NS)

EB powder was purchased from Kremer Pigmente GmbH & Co. KG. EB-NS and prepared as follows. EB powder was ground with a ceramic pestle and mortar to mechanically reduce the crystal size until the powder was visibly brighter, a consequence of the dichroism of EB^25^. The ground powder (10 mg) was transferred to a glass vial together with 5 mL of isopropanol (Fischer Chemical, 99.98%). Tip sonication was performed in a glass vial with a Fisherbrand™ Model 120 Sonic Dismembrator (Fischer Scientific) in an ice bath for 1-6 h at 60-72 W. For nIR imaging, bleaching experiments and MSD analysis, the mixture was allowed to settle for 5 min before tip sonication. The supernatant was then decanted into another glass vial and sonicated. EB dispersions were stored at room temperature and vortexed prior to use (Vortex Mixer VV3, VWR International) for 10 s at maximum power (2500 min^−1^, 10 W).

### Atomic Force Microscopy (AFM)

100 μL of EB-NS stock dispersion (6 h-sonicated) was diluted 10x with isopropanol and vortexed for 10 s. 10 μL of EB-NS were spin-coated (G3 Spin Coater, Specialty Coating Systems, Inc.) on a mica surface. The substrate was kept spinning at 500 RPM (7 RCF) for 2 min (5 s of ramp time, 30 s of dwell time). An Asylum Research MFP-3D Infinity AFM (Oxford Instruments) was employed in AC mode (software version 15.01.103). Rectangular cantilevers from Opus (160AC-NA, MikroMasch Europe) were used (aluminum coating, tetrahedral tip, 300 kHz resonance frequency, force constant of 26 N·m^−1^). Image analysis was performed with Gwyddion (version 2.51).

### nIR Fluorescence Spectroscopy

The setup consists of a monochromator (MSH-150, LOT-Quantum Design GmbH) equipped with a xenon arc lamp and a diffraction grating, an Olympus IX73 microscope with a 10x objective (UplanFLN 10x/0.30, Olympus), and a Shamrock 193i spectrograph (Andor Technology Ltd.) coupled to an array nIR detector (Andor iDUs InGaAs 491). Spectra were recorded from EB-NS dispersions at an excitation wavelength of 615 nm with an exposure time up to 5 s and a slit width up to 500 μm. The Andor SOLIS software (version 4.29.30012.0) was employed for spectra acquisition and further analyzed with OriginPro 8.1. For 2D spectra, EB-NS were placed on glass substrates. The excitation wavelength was scanned in 4 nm steps with the monochromator, and at each wavelength a spectrum was recorded with an integration time of 1 s at a slit width of 10 μm. The 2D spectra were corrected for the quantum efficiency of the detector as well as the spectral irradiance of the xenon lamp of the monochromator using a self-written Python script.

### Fluorescence Saturation Measurements

The main components of the employed setup were a laser source (Supercontinuum laser SC400-4-20, Fianium), a photodetector (single photon avalanche diode PDM series, MPD) and a 60x objective lens (Apo N, 60x/1.49 NA oil immersion, Olympus). 10 μL of the supernatant of a 6 h-sonicated EB-NS sample (1:100 diluted in isopropanol) were spin-coated on a glass cover slide. Spin-coating parameters were the same ones chosen for AFM measurements. Despite the polydispersity of EB-NS, scanning them using a confocal microscope through the diffraction limited focal spot of 1.49 NA objective allowed us to select only the smallest particles with sizes estimated to be not exceeding ≈ 100 nm. The spot size of larger EB-NS exceeded the dimensions of a diffraction limited focal spot, thus allowing us to distinguish them from smaller particles. After the dimmest particles could be detected on the sample surface, we acquired its fluorescence images at different excitation powers. The dependence between the brightness of the EB-NS and the excitation light power (measured directly before the objective lens) was studied. The excitation wavelength was 640 nm.

### Fluorescence Polarization Measurements

The setup mainly consisted of a light source (LED, Lumencor) which emitted unpolarized excitation light, a photodetector (EMCCD camera, iXon Ultra DU-897U-CS0, Andor) and a 60x objective lens (Apo N, 60x/1.49 NA oil immersion, Olympus). Measurements were performed by recording fluorescence images of individual EB-NS at different positions on a linear polarizer, which was placed in front of the camera. The sample preparation steps coincided with the ones performed for fluorescence saturation measurements. Atto 488, purchased from Atto-Tec, was used as a reference.

### nIR Fluorescence Imaging Setup

An Olympus BX53 microscope equipped with 20x (MPlanFL N 20x/0.45, Olympus) and 100x (UPlanSApo 100x/1.35 Sil, Olympus) objectives was used. A Xeva-1.7-320 nIR camera (Xenics®) and a Zyla 5.5 sCMOS camera (Oxford Instruments) were used to observe the EB-NS fluorescence excited by a 561 nm laser (Cobolt Jive™ 561 nm). Typically, a droplet of an EB-NS dispersion was put on glass slides, dried and imaged at 10-50 mW excitation power. To assess bleaching, a solution of Rhodamin B (Sigma-Aldrich) in isopropanol was prepared, and a drop of this solution was put on a glass slide for imaging. Dried EB-NS were put on a separate glass slide. At a continuous excitation of 100 mW at 561 nm, images with an integration time of 100 ms were recorded 120 times every 0.5 min in the case of Rhodamin B, and every 1 min with an integration time of 1 s in the case of EB-NS. For Rhodamin B the Zyla camera and for EB the Xenics nIR camera was used. The specified fluorescence intensity corresponds to the mean gray value of the images.

### Microrheology and Correlative Size/Intensity Measurements

For microrheology measurements, a 2 mL centrifuge tube was charged with 1 mg of dried EB-NS and 0.5 mL of glycerol (Alfa Aesar, 99+%). The powder was re-dispersed in glycerol by 2 min of tip sonication in an ice bath at 60 W power. This solution was then centrifuged for 5 min at 15000 RCF in order to remove larger sheets from the dispersion. A drop of this solution was put on a glass slide for imaging using the Xenics camera at 100 mW excitation with a Cobolt 561 nm laser. Movies consisting of 1000 frames in 1 s intervals with an integration time of 1 s were recorded and analyzed in ImageJ (1.52i) with the Mosaic Particle Tracker^61^. The following parameters were employed: radius = 4, cutoff = 0.001, per/abs = 1.5, link range = 3, displacement = 1, dynamics = Brownian. Trajectories with a length smaller than 50 frames were not considered in the analysis. Additionally, trajectories with nonlinear MSD plots due to stage movements etc. were filtered out. The fluorescence intensity specified in fig. 4g is equal to the zeroth moment of the intensity distribution function as determined by the custom-written tracking algorithm.

The Stokes radii were calculated the following way. The tracking of the Brownian motion of an EB-NS yields the trajectory of the particle as a set of time-dependent *x* and *y* positions for *N* time steps of length *τ*. Example trajectories are shown in fig. 4e. From the *x* and *y* positions, the square displacement for the *n*-th time step 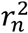 is calculated as 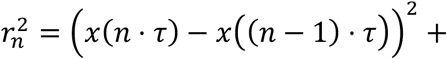 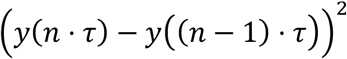, where *n* ranges from 1 to *N*. The mean square displacement 〈*r*^2^〉 (MSD) is simply the mean over all single time step values: 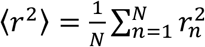. Assuming the diffusion is restricted to two dimensions, the diffusion coefficient *D* and the MSD are linked *via* the relation 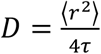. Finally, from the Stokes-Einstein equation, the Stokes radius *R* is calculated as 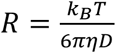, where *η* is the dynamic viscosity of the solvent, *T* the temperature and *k*_B_ the Boltzmann constant. Here, a temperature of 293 K and a dynamic viscosity of glycerol of 1412 centipoises were assumed^62^. The Stokes-Einstein equation is only strictly valid for spherical particles. Due to the high anisotropy of EB-NS, we assumed that Brownian motion is dominated by the diameter of the nanosheets and not the much smaller height.

### Scanning Electron Microscopy (SEM)

A 6 h-tip sonicated sample was observed under a LEO SUPRA 35 microscope (Zeiss) with an Inlens detector at 20 kV (secondary electrons). 10 μL of EB-NS suspension were drop-casted on a Si wafer. Both gold sputtering and evaporation (≈ 2 nm of gold layer) were tested on different samples, nevertheless the best imaging conditions were met when no gold deposition step was performed at all. Interestingly, EB was conductive enough to be seen at SEM without gold deposition.

### *In vivo* Microrheology of *Drosophila* Embryos

*Drosophila* embryos expressing Histone2Av-GFP^63^ with an age of 0-1 hour were collected and dechorionated with hypochlorite for 120 s, washed with water thoroughly, aligned on a piece of agar, transferred to a cover slip coated with glue and covered with halocarbon oil (Voltalef 10S, Lehmann & Voss) after a slight desiccation. An aliquote of suspension in water of a 6 h-tip sonicated EB-NS sample (filtered with a 0.20 μm syringe filter) was injected using Microinjector FemtoJet® (Eppendorf) on an inverted microscope: the injection volume is calculated according to literature^64^. After around 30 min of incubation, the sample could be moved to the previously mentioned nIR setup for co-localization experiments. EB-NS were excited with the 561 nm laser (300 mW), while a fluorescence lamp (X-Cite® 120Q, Excelitas Technologies) was used for GFP excitation. Both channels were observed through a 100x objective and recorded with a Zyla 5.5 sCMOS camera. An exposure time of 0.1 s was chosen and images on both channels were taken for 60 s (10 fps). A 2×2 pixel binning was performed to lower the size of acquired data and thus facilitate the following particle tracking analysis. As for the EB-NS tracking in glycerol, ImageJ with the Mosaic Particle Tracker was employed to analyze the acquired frames. Only the EB data was evaluated, since the nuclei did not display any significant motion during the acquisition. The following parameters were employed for analysis: radius = 5, cutoff = 0, per/abs = 0.1, link range = 100, displacement = 2, dynamics = Brownian. The diffusion coefficient was then evaluated as previously described. The radius of each EB-NS was estimated from 10 images of the movie. Finally, the Stokes-Einstein equation was used to estimate a corresponding dynamic viscosity.

### Stand-off Detection

nIR images were acquired with a CMOS based DCC3240M camera (Thorlabs), equipped with a 900 nm (FEL0900, Thorlabs) and a 920 nm (LP920, Midwest Optical Systems) longpass-filter, mounted in series in order to exclude visible background. A white light source (UHP, Prizmatix) connected to a 700 nm (FESH0700, Thorlabs) and a 750 nm (FESH0750, Thorlabs) shortpass-filter was used for excitation. Exposure times between 50-100 ms were used for all acquisitions. ssDNA modified SWCNTs were obtained by placing 125 μl (2 mg/mL in PBS) (AT)_15_ ssDNA (Sigma Aldrich) and 125 μl (6,5) chirality enriched SWCNTs (Sigma Aldrich, Product No. 773735) (2 mg/mL in PBS) for tip sonication (15 min, 30% amplitude). The obtained suspension was centrifuged twice for 30 min at ambient temperature (16100 g). The amounts of ICG (≈ 90%, MP Biomedicals GmbH) and of a 6 h-tip sonicated sample of EB-NS in water were ≈ 0.1 mg/mL.

Seeds of *Arabidopsis thaliana* (Col.0 ecotype) were sown on sterilized soil (8 h, 80 °C) and stratified for 2 days in the dark at 4 °C. The plants were grown under long day-length (16 h light/8 h dark) in climate controlled growth chambers at 22 °C, 60% humidity and light intensity of 120-150 μmol m^−2^ s^−1^. Delivery of nanomaterials and buffer into leaves of *Arabidopsis* was performed as described by Giraldo *et al*^51^. In brief, 50 μl of the desired nIR fluorophore suspension were infused through the lower (abaxial) side of the leaf lamina with a needle-less syringe. Spectroscopic measurements for the presence of EB-NS in the leaf was performed using the nIR fluorescence spectroscopy setup (5 s exposure time) as described above. Regarding the imaging of EB-NS under room light excitation, 1 mL of supernatant taken from a 6 h-sonicated sample in water was observed. A background picture was taken and used for background subtraction.

## ASSOCIATED CONTENT

Supporting Information. Supporting Information is available for this article.

## DATA AVAILABILITY

The data that support the findings of this study are available from the corresponding author upon reasonable request.

## CODE AVAILABILITY

The codes that support the findings of this study are available from the corresponding author upon reasonable request.

## AUTHOR INFORMATION

Corresponding Author

## ACKNOWLEDGMENT

We thank the Volkswagen foundation and the life@nano cluster for funding (S.K.). We thank Dr. Burkhard Schmidt for recording reflectance spectra, Elena Polo for initial exfoliation experiments, Nelli Teske and Jeremias Sibold for help with the SEM and Angela Rübeling for technical assistance. Furthermore, we thank Wentao Peng/Prof. Dr. Vana for assistance with zeta potential measurements. We thank the Steinem and Janshoff labs for discussions and support. We are grateful to Dr. Ellen Hornung for providing us with *Arabidopsis* plants and related information. This work was funded by the National Science Foundation under Grant No. 1817363 to J.P.G.

## AUTHOR CONTRIBUTIONS

S.K. conceived and designed the study. G.S. performed AFM experiments. G.S., H.P. and A.S. exfoliated materials and collected spectra. H.P. performed and analyzed correlative experiments. G.S. together with R.N., F.A.M, Z.L. performed biological imaging experiments with inputs from L.E., J.P.G. and J.G. A.C. performed single photon counting and anisotropy experiments. T.A.O. performed cell viability experiments. G.S., H.P., J.P.G., A.C. and S.K. analyzed data and wrote the manuscript with contributions from all authors.

