## Supplementary Information for "Exfoliated near infrared fluorescent CaCuSi_4_O_10_ nanosheets with ultra-high photostability and brightness for biological imaging"

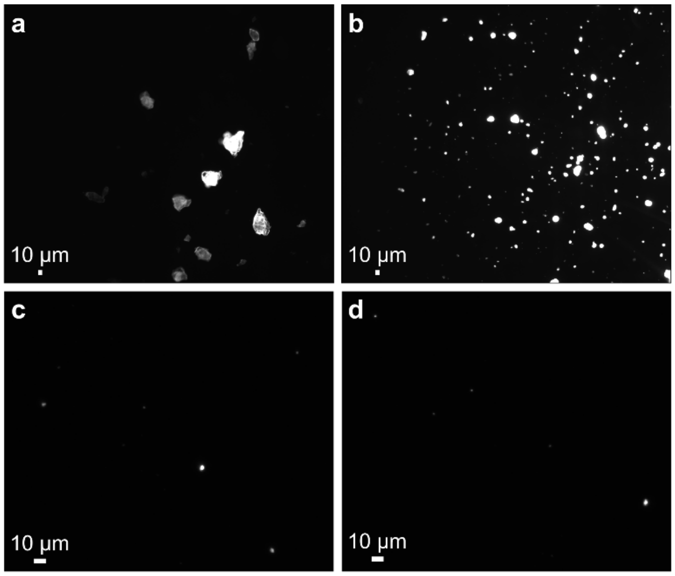


***Fig. S1: nIR fluorescence images of Egyptian Blue (EB) before and after tip sonication. a*** *EB bulk powder dispersed in isopropanol.* ***b*** *Egyptian Blue nanosheets (EB-NS) after 6 h-tip sonication.* ***c,d*** *EB-NS after 6 h-tip sonication and size-cut-off filtration (d = 0.45 µm). All samples were drop-casted (10 µL) on glass cover slides before imaging. Every purification step reduces the overall concentration of particles but increases monodispersity. Note that the contrast was not adjusted. Therefore, the smaller EB-NS in b-d are difficult to see (compared to the larger ones) and it looks like there are not many particles left but when zooming in they are visible.*

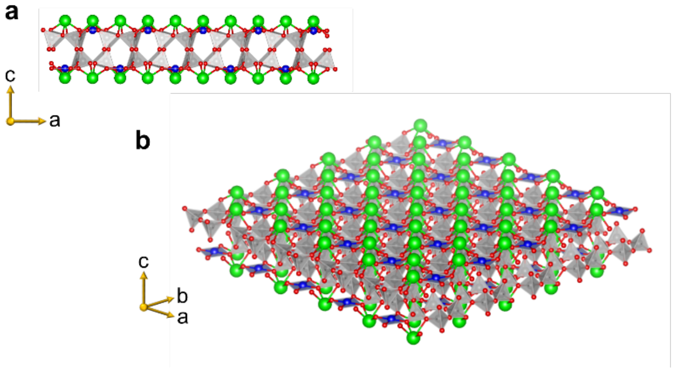


***Fig. S2: Schematic of a monolayer of EB. a*** *Frontal view.* ***b*** *3D axonometric projection. For both illustrations, EB neutron powder diffraction data obtained from literature was used[1]. Si, O, Ca and Cu atoms are depicted as gray, red, green and blue spheres, respectively.*

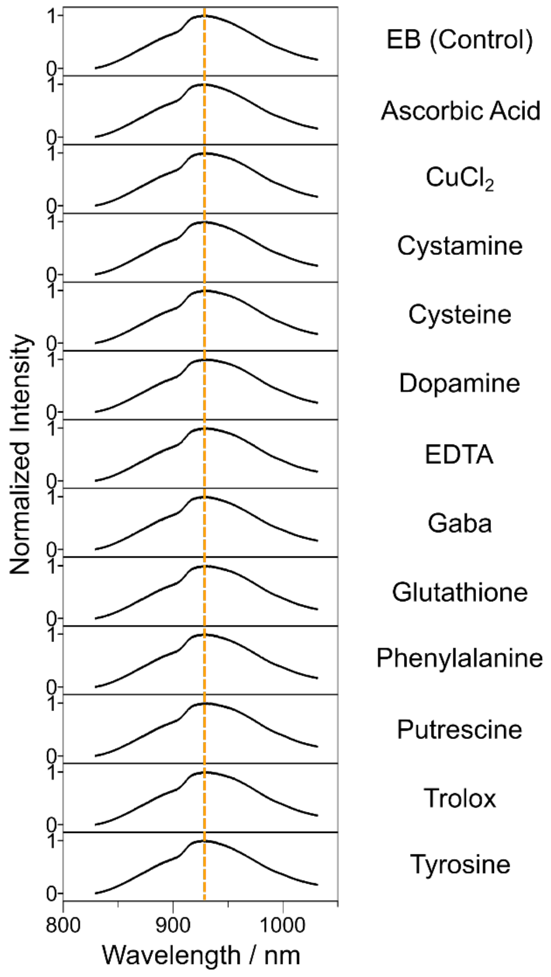


***Fig. S3: Fluorescence response of EB-NS to different analytes of interest.*** *nIR spectra of EB-NS (6 h sonication, 10 µg/mL) in water 10 min after addition of analytes (100 µM) that are known to affect spectra of other fluorophores. The EB-NS fluorescence did not display any significant shifts. This result further shows the stable nIR fluorescence of this nanomaterial.*

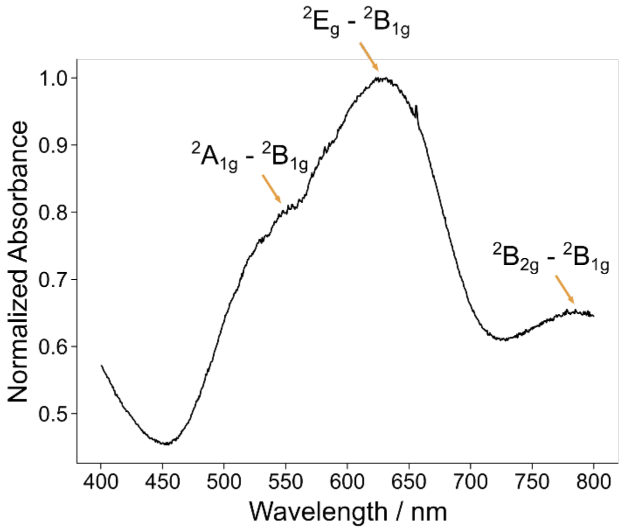


***Fig. S4: Absorption (reflection) spectrum of EB powder.*** *Three broad bands corresponding to different electronic transitions are observed (yellow arrows). The symmetry species of the orbitals involved in the transition are indicated next to the arrows. The attribution of the bands to the symmetry species was made according to the model proposed by Accorsi et al.*[2]*.*

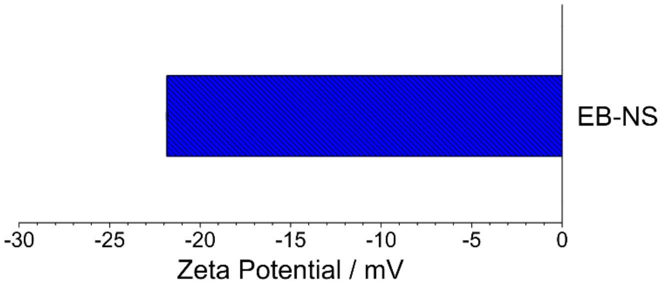


***Fig. S5: Zeta potential of EB-NS.*** *Zeta potential of EB-NS in water (2 mg/mL). Error bars correspond to the standard deviation of triplicates.*

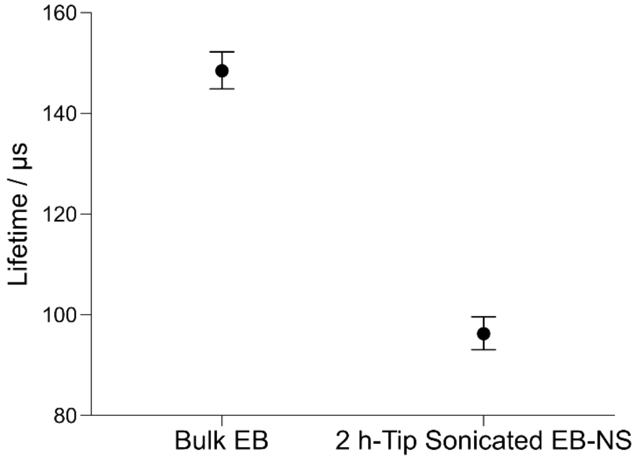


***Fig. S6: Fluorescence lifetimes of bulk EB and exfoliated EB-NS.*** *Error bars correspond to standard deviations (n = 100 repetitions performed for each sample).*

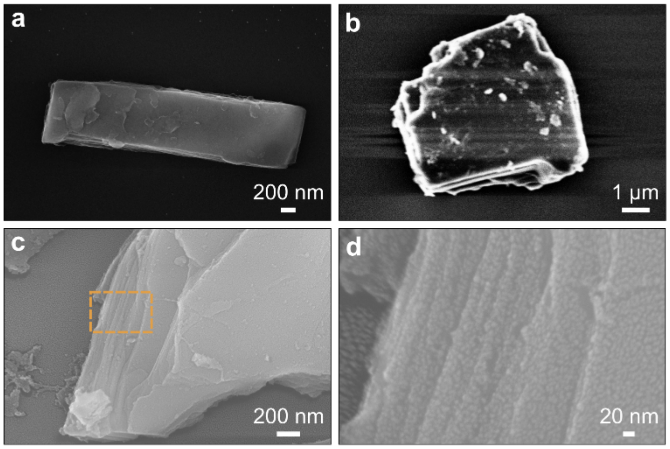


***Fig. S7: Scanning electron microscopy (SEM) images of larger exfoliated EB (nano)sheets.*** *Different samples and preparation techniques are shown.* ***a*** *No gold deposition.* ***b*** *≈ 2 nm evaporated gold.* ***c*** *≈ 2 nm of sputtered gold on the surface of the sample.* ***d*** *A magnified region from the inset shown in c indicates the layered structure.*

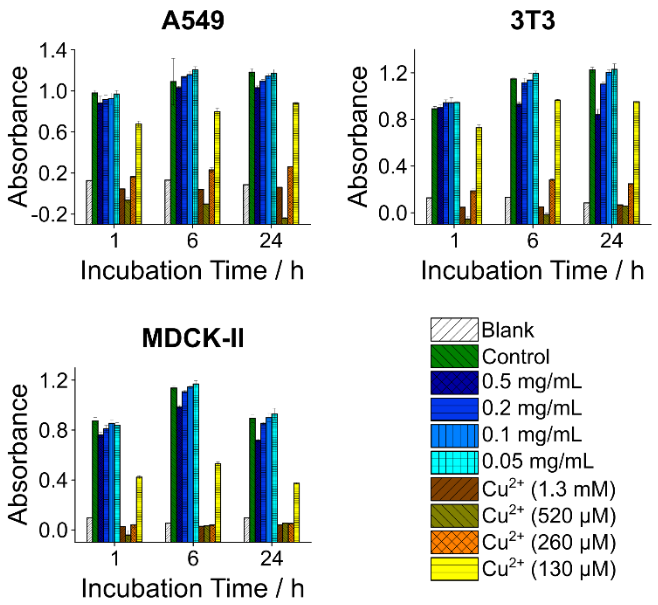


***Fig. S8: Cell viability in the presence of EB-NS.*** *Cytotoxicity of EB-NS was assessed for 3 different common cell lines (A549, NIH 3T3 and MDCK-II) using a standard assay (see materials and methods). EB-NS did not show significant effects on cell viability, which proves this material's biocompatibility. As a control we used* $Cu^{2+}$ *ions (*$CuSO_4$*), which drastically decreased viability. N = 4 independent samples (quadruplicates) for each data point, error bars correspond to standard deviation.*

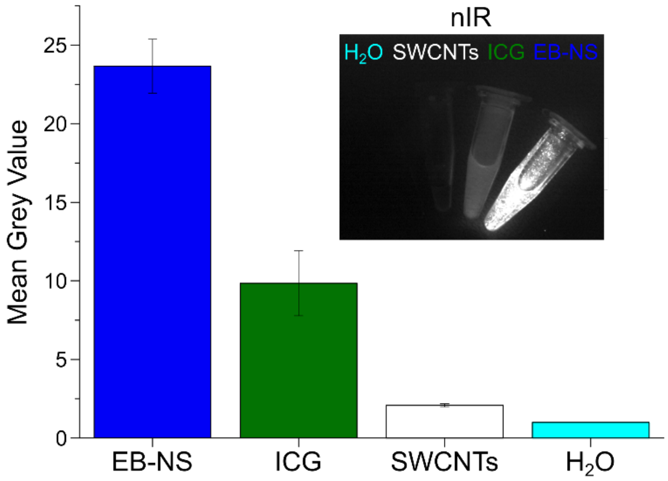


***Fig. S9: Relative fluorescence intensity of different nIR fluorophores in the low-cost stand-off detection setup.*** *Average total emission intensities (i.e. mean grey pixel values) of three different nIR fluorophores in Eppendorf tubes at a similar concentration (≈ 0.1 mg/mL). Data were normalized to the water control. EB-NS, ICG and SWCNTs respectively yielded mean grey values of 23.7 ± 1.7, 9.8 ± 2.1 and 2.1 ± 0.1 (mean ± standard deviation). N = 3 independent samples.*

### SUPPLEMENTARY MATERIALS AND METHODS

#### Absorption (reflection) spectrum of Egyptian Blue (EB) powder

Absorption spectra were recorded with an AvaSpec-UV/VIS/NIR two channel broad band spectrograph (Avantes) containing a balanced deuterium-halogen lamp (AVALIGHT-DH-S-BAL). Only the signal recorded in the UV/Vis channel is shown, which is equipped with a 2048L UV/VIS spectrometer with a 25 µm slit and a 300 lines/mm grid. The Avasoft-Full software was used for recording.

#### Cytotoxicity Tests of EB-NS

A549, NIH 3T3 and MDCK II cell lines were employed in this study.

- A549 cell line and NIH 3T3 cell line: $D10F^+$ (DMEM +4.5 g/L glucose with L-glutamin (4 mM), 10% fetal bovine serum and 100 µg/mL Pen-Strep), trypsin/EDTA (0.05/0.05%) solution;
- MDCK II cell line: $M10F^+$ (EMEM with Earle's salts with L-glutamin (2 mM), 10% fetal bovine serum and 100 µg/mL Pen-Strep), trypsin/EDTA (0.25/0.05%) solution.

The cell viability (MTS) assay was performed using CellTiter 96® AQueous One Solution Cell Proliferation Assay (Promega, G3580). 2 mL (≈ 4 mg) of a 6 h-sonicated sample of EB-NS in water (2 mg/mL) was centrifuged (10 min, 13100 RCF). The supernatant was removed and the so-obtained pellet was re-dispersed in 400 µL, in order to reach a concentration of ≈ 10 mg/mL of EB-NS. The sample was diluted to 5% (0.5 mg/mL), 2% (0.2 mg/mL), 1% (0.1 mg/mL) and 0.5% (0.05 mg/mL).

A 13 mg/mL (i.e. 52 mM) water solution of $CuSO_4{\cdot}5H_2O$ (98%, J&K Scientific) was used as a reference to assess the maximum sensitivity of the cells to $Cu^{2+}$ ions. 0.2% (100 µM) corresponds to the copper amount in 2% of the 2 mg/mL EB-NS dispersion (i.e. 0.04 mg/mL of EB-NS). Therefore 2.5% (1.3 mM), 1% (520 µM), 0.5% (260 µM) and 0.25% (130 µM) correspond to the EB-NS concentrations of 0.5 mg/mL, 0.2 mg/mL, 0.1 mg/mL, 0.05 mg/mL, respectively. The $Cu^{2+}$ concentrations for this experiment have been shown to cause varying degrees of cytotoxicity in literature[3]. 2% (v/v) deionized water was employed as positive control. Cells were initially incubated in a 96-well plate at $1.2\times10^3$ cells per well for 24 h in $M10F^-$ Media at 37 °C and 7.5% $CO_2$ (MDCK II), or in D10F-Media at 37 °C and 5% $CO_2$ (A549 and NIH 3T3). A suspension of EB-NS was diluted to the different concentrations in appropriate media and added to the cells to be incubated for 1, 6 or 24 h. Viability of the cell samples was determined by MTS-assay.

**Fluorescence Lifetime Measurements**

Lifetime measurements (frequency domain) were conducted in a well-plate using a Firesting oxygen meter from Pyroscience (Aachen, Germany) at room temperature. The excitation wavelength was set to 620 nm. An excitation frequency of 4 kHz and an LED intensity of 40% were employed.

**Zeta Potential Measurements**

A Zetasizer Nano S device (Malvern Instruments) was employed for this experiment. The obtained dataset was analysed *via* the Zetasizer software.
